# signifinder enables the identification of tumor cell states and cancer expression signatures in bulk, single-cell and spatial transcriptomic data

**DOI:** 10.1101/2023.03.07.530940

**Authors:** Stefania Pirrotta, Laura Masatti, Anna Corrà, Fabiola Pedrini, Giovanni Esposito, Paolo Martini, Davide Risso, Chiara Romualdi, Enrica Calura

## Abstract

Over the last decade, many studies and some clinical trials have proposed gene expression signatures as a valuable tool for understanding cancer mechanisms, defining subtypes, monitoring patient prognosis, and therapy efficacy. However, technical and biological concerns about reproducibility have been raised. Technical reproducibility is a major concern: we currently lack a computational implementation of the proposed signatures, which would provide detailed signature definition and assure reproducibility, dissemination, and usability of the classifier. Another concern regards intratumor heterogeneity, which has never been addressed when studying these types of biomarkers using bulk transcriptomics.

With the aim of providing a tool able to improve the reproducibility and usability of gene expression signatures, we propose *signifinder*, an R package that provides the infrastructure to collect, implement, and compare expression-based signatures from cancer literature. The included signatures cover a wide range of biological processes from metabolism and programmed cell death, to morphological changes, such as quantification of epithelial or mesenchymal-like status. Collected signatures can score tumor cell characteristics, such as the predicted response to therapy or the survival association, and can quantify microenvironmental information, including hypoxia and immune response activity.

*signifinder* has been used to characterize tumor samples and to investigate intra-tumor heterogeneity, extending its application to single-cell and spatial transcriptomic data. Through these higher-resolution technologies, it has become increasingly apparent that the single-sample score assessment obtained by transcriptional signatures is conditioned by the phenotypic and genetic intratumor heterogeneity of tumor masses. Since the characteristics of the most abundant cell type or clone might not necessarily predict the properties of mixed populations, signature prediction efficacy is lowered, thus impeding effective clinical diagnostics. Through *signifinder*, we offer general principles for interpreting and comparing transcriptional signatures, as well as suggestions for additional signatures that would allow for more complete and robust data inferences. We consider *signifinder* a useful tool to pave the way for reproducibility and comparison of transcriptional signatures in oncology.

## Introduction

Intratumor heterogeneity refers to the diversity of the gene mutation spectrum and the biological characteristics between the tumor and its microenvironment, found in different parts of the same mass. Tumor heterogeneity can impact treatment efficacy and is important to address for the understanding of tumor dynamics^1^. The past ten years have seen increasing advances in the resolution of cancer transcriptome detection through the use of single-cell RNA sequencing (scRNA-seq) and spatial transcriptomics. These new technologies are providing a growing body of evidence that recognizes cancers as cell mosaics, with highly heterogeneous behavior that is often guided by spatial patterns created by clonal cells and local microenvironmental stimuli^2–4^. To address this complexity, computational methods are needed to enable the characterization of tissue heterogeneity^5^ using the vast amount of information gained by high-resolution transcriptomic technologies.

An emerging trend in single-cell and spatial transcriptomic analyses is the identification of cancer cell states. This involves recurrent gene-expression programs that define neoplastic and tumor microenvironment (TME) cell behaviors, thus dissecting tumor complexity. Cancer cell states are defined through the use of gene expression modules that have been summarized into cell-specific scores^4,6–10^. Despite some promising early results, we are far from having a complete list of relevant gene modules for all cancers. Such a list would be a key tool for genomic data interpretation, however, even if a comprehensive set of modules were available, they would likely miss information on inter-patient variability levels, since their identification is currently based on the analysis of relatively few samples.

On the other hand, after decades of bulk gene expression studies on large patient cohorts, several transcriptional signatures are available that seem to be good indicators of cancer phenotypes^11^. Similar to cancer cell states, transcriptional signatures are based on the expression of a specific gene set (sometimes including coefficients that weigh the gene contributions), summarized into a score that provides single-sample predictions. Transcriptional signatures have been frequently studied because of their potential to show ongoing cancer activities. This feature could help us guide therapeutic decisions and monitor interventions, understand cancer mechanisms, define tumor subtypes, and assess patient diagnosis and prognosis^12,13^. Transcriptional signatures are also used to assess the complex relations between the tumor and its normal counterpart, called the TME. This has been shown to not only confound data analysis (the heterogeneous mixture of cell types affects tumor purity and can bias tumor data analysis) but also to be an intrinsic tumor attribute, which is worth considering for sample characterization (e.g., signatures to monitor intrinsic and acquired immune resistance^14^. Since it has been demonstrated that bulk-based functional analysis tools that use manually curated footprint gene sets can be applied to scRNA-seq data^15^, we thought that the plethora of bulk-derived transcriptional signatures could be also used to dissect the complexity of high-resolution transcriptomics.

However, even after decades of research, the field of bulk transcriptional signatures still has issues to solve. Firstly, though many gene expression-based prognostic signatures have been reported in the literature, very few are used in clinical practice^16^. The way to achieve accurate tumor classifications based on transcriptional signatures is debated and furthermore complicated by the lack of standard practices^12,17^. Moreover, their reproducibility and dissemination are affected by the lack of public open-source implementations: the vast majority of signatures are not published along with their computational code and only a few of them have been implemented in software packages.

Thus, with the intent of using bulk transcriptional signatures to define cancer cell states in single-cell and spatial transcriptomics, we developed the R/Bioconductor package *signifinder*. This serves as a bridge between signature discovery and signature usability in all three types of transcriptomic data: bulk, single-cell, and spatial transcriptomics. To accomplish this task, *signifinder* provides the infrastructure to collect and implement cancer transcriptional signatures that are available in the literature. This software has been conceived as an opensource R package built around the gene expression data structures of the Bioconductor project. This framework guarantees the interoperability of *signifinder* with most of the bulk, singlecell, and spatial transcriptomics data analysis workflows, also paving the way for a systematic examination and comparison of the available signatures.

With *signifinder* we can assess, for the first time, the extent to which bulk transcriptional signatures behave in intra-masses when applied to single-cell or single-spot spatial transcriptomes. Here, we present three case studies, with in-depth analysis and discussion about the use of transcriptional signatures in high-resolution transcriptomic datasets. These findings provide intriguing evidence on the nature and extent of the use of cell-resolution transcriptional signatures in oncology, potentially leading to new research directions.

## Materials and Methods

### TCGA ovarian cancer RNA-seq data preparation

We downloaded the The Cancer Genome Atlas (TCGA) ovarian cancer (OVC) *RNASeqGene* data from the *curatedTCGAData* package^18^ (version 2.0.1), a *SummarizedExperiment* object with the raw-count gene expression values. Then, data were normalized using the *betweenLaneNormalization* function from the *EDASeq* package^19^ (version 2.32.0, *which = “median”)*. After that, we computed all available signatures for ovary and pan-tissue cancers that were collected within *signifinder*. We computed the correlation matrix of the signature scores and performed a hierarchical clustering to group similar correlations. For this step we used the *dist* function (*method* = “*euclidean*”) and the *hclust* function (*method* = “*complete*”) from the *stats* package (version 4.2.2). We then computed the silhouette index to find the best number of clusters and decided to discuss the ones containing the higher correlation values (*NbClust* package^20^, version 3.0.1). Signature score plots were made with *signifider* and *ggplot2* packages^21^ (version 3.4.0).

### Single-cell glioblastoma data preparation

We downloaded the glioblastoma (GB) scRNA-seq dataset of Darmanis et al. (Gene Expression Omnibus ID: GSE84465^22^) using the *TMExplorer* R package (version 1.6.2^23^). The *SingleCellExperiment* object provides the raw count expression values. We selected two samples, one containing the core cells and the other the peripheral cells of the tumor mass from the BT_S2 patient, and we removed the zero-expression genes. Then, using *scone* workflow^24^ (version 1.22.0), we applied the best suggested normalization method (method = “none, fq, ruv_k=4, no_bio, no_batch”, which stands for no data imputation, full-quantile normalization, four RUVg factors, no biological or batch information to consider). Additionally, to reduce the large number of false zeros, we ran ALRA on the normalized dataset^25^. Then, to have sizable cell groups for signature score comparisons, we kept only the cell types that had greater than 20 cells present in both core and peripheral samples (Additional Table S1, cell type annotations by Darmanis et al. were used). Immune cells, neoplastic, and oligodendrocyte precursor cells were considered for further analysis (1054 cells). We performed t-distributed stochastic neighbor embedding (t-SNE) using the top 50 components of the principal component analysis (*scater* package^26^, version 1.26.1). Finally, we computed all the signatures for brain and pantissue cancers that are available in *signifinder*. Differences between signature scores in different cell types were computed using *t test* statistics. Plots were made using *signifinder, scater*, and *ggplot2* R packages.

### Spatial Transcriptomics Breast Cancer data preparation

We downloaded the spatial transcriptomic dataset “Human Breast Cancer: Ductal Carcinoma In Situ, Invasive Carcinoma (FFPE)”, included in the 10x Genomics Visium Spatial Gene Expression data, from the 10x website (https://www.10xgenomics.com). This dataset was generated following the *10x Visium* spatial gene expression protocol for Formalin-Fixed Paraffin-Embedded (FFPE) specimens, the library was sequenced using an Illumina NovaSeq sequencer, and processed and quantified using Space Ranger software (version 1.3.0; 10x Genomics proprietary software). Annotation of the histology image was performed by an expert anatomopathologist, and spots were annotated using the Loupe Browser interface (version 3.0, 10x Genomics). Then, we imported the spot gene expressions and their annotations in the form of a *SpatialExperiment* using the *read10xVisium* function from the *SpatialExperiment* package^27^ (version 1.8.0) and we normalized the raw counts using the *logNormCounts* function from the *scater* R package^26^ (version 1.24.0). Then, we computed all the signatures for breast and pan-tissue cancers available in *signifinder*. Plots were made with *signifinder* and *ggspavis* R packages (version 1.4.0).

## Results and Discussion

### Design and implementation

Figure 1 graphically outlines *signifinder* development and the analysis workflow. The package contains a collection of signatures from the literature, together with their implementations (Figure 1A). As the input, the user can provide transcriptional bulk data of cancer samples (either from microarray or RNA-seq technologies), single-cell sequencing data, or spatial transcriptomics data, and the package returns signature scores at the level of samples, cells, or spots, respectively (Figure 1B). Finally, *signifinder* can provide graphical summaries that are helpful to visually represent single signatures or compare multiple signatures (Figure 1C).

**Figure 1.**
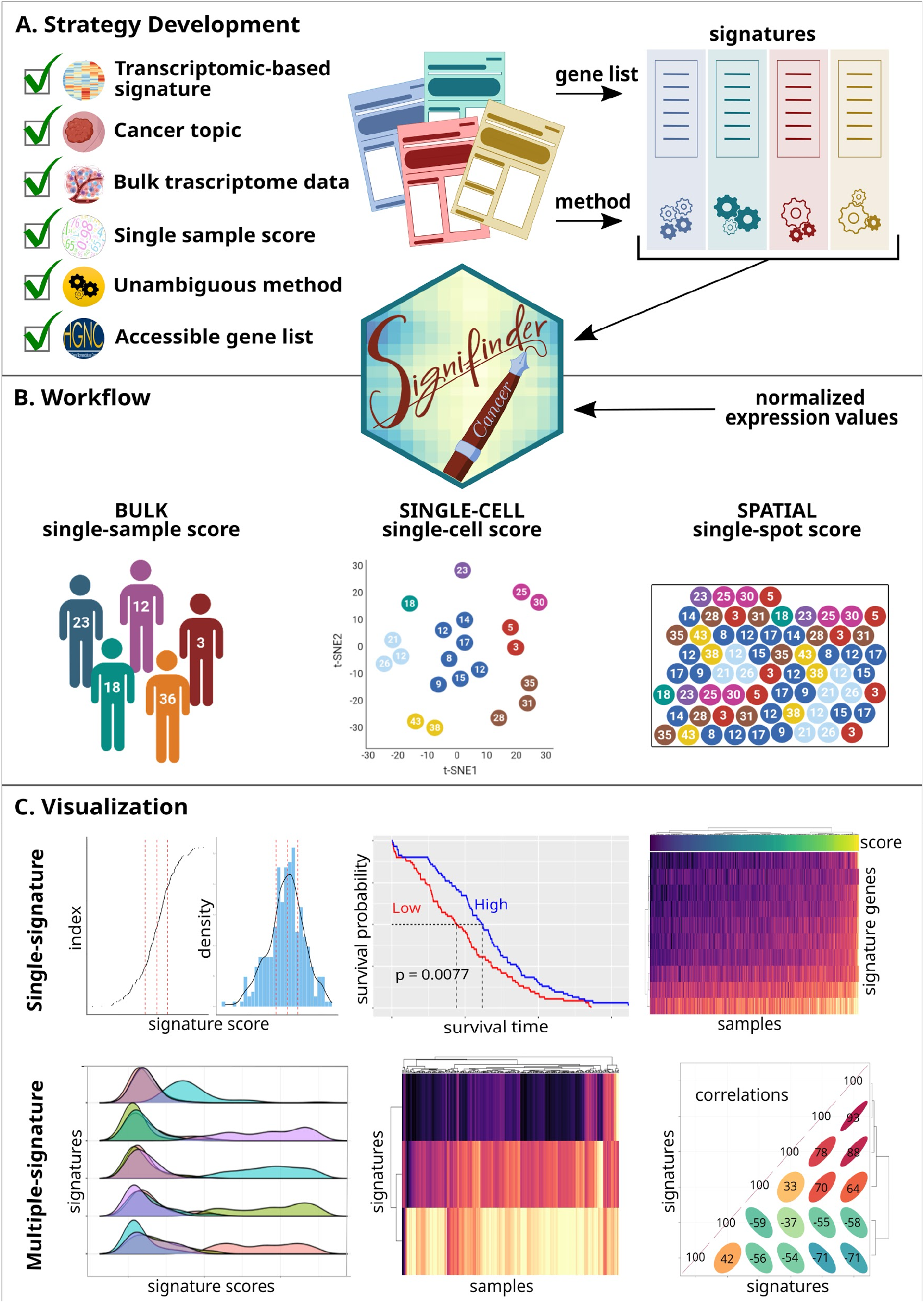
signifinder implementation and workflow. **A**) The scheme for signifinder development: following stringent criteria, we collected and implemented the lists of genes and algorithms for signature computations. **B)** signifinder workflow starts with gene expression data from bulk, single-cell, or spatial transcriptomics and, without any other type of prior knowledge, the user can retrieve and compare single sample, cell, or spot scores, respectively. **C)** signfinder offers several plots to explore and compare the signature score distributions.

### The collection of signatures

We established a set of stringent criteria for the inclusion of signatures: *(i)* signatures should rely on cancer topics, and be developed and used on cancer samples; *(ii)* signatures should exclusively use transcriptomic data, though exceptions have been made in case of combination of gene expression and signature-related gene weights; *(iii)* signatures must release a clear gene list used for the signature definition, where all genes have an official gene symbol (Hugo consortium) or an unambiguous translation (genes without an official gene symbol are removed); *(iv)* the method to calculate expression-based scores should be unambiguously described; *(v)* additional clarity about the type of expression in the input (e.g., counts, log counts, FPKM, or others) may also be required.

The first step was a literature search using the following keywords: “cancer”, “gene expression”, “microarray” or “RNA sequencing”, and “signature”, providing an initial set of 2000 journal articles. We then excluded papers on mutational signatures and those including microRNAs or other omic features such as DNA methylations, which accounted for a large part of the considered articles. Then, we focused on articles that proposed a patient-specific summary score. We ended up with 150 papers that were screened manually, applying the above selected criteria. In the end, the selected signatures cover 16 cancer topics (including alteration in the epithelial-to-mesenchymal transition (EMT) process, chromosomal instability (CIN), angiogenesis, hypoxia, altered metabolism, and cell cycle rate; see Table 1 for the complete list). Some signatures are dedicated to the study of tumor cell composition and its relationship with the microenvironment, i.e., cancer stem cell presence, signatures for monitoring immune system activity, extracellular matrix (ECM) composition, and angiogenesis activity. Finally, some signatures monitor clinical outcomes such as chemo-resistance or patient prognosis. While it may not ever be possible to include all cancer signatures proposed in the literature, our package makes the addition of new signatures (by us or by others via “pull requests”) extremely easy. Moreover, in the current release of *signifinder*, all the included signatures rely on bulk tumor expression experiments, even if the package infrastructure could potentially store and manage signatures derived by single-cell and spatial transcriptomics. As stated in the introduction, some of these high-resolution signatures are starting to appear but they lack performance evaluations at the inter-patient variability level, since their identification is currently based on the analysis of relatively few samples. The interchangeability of signatures across transcriptional omics, as is proposed by *signifinder*, would improve signature evaluation and applicability.

**Table 1.**
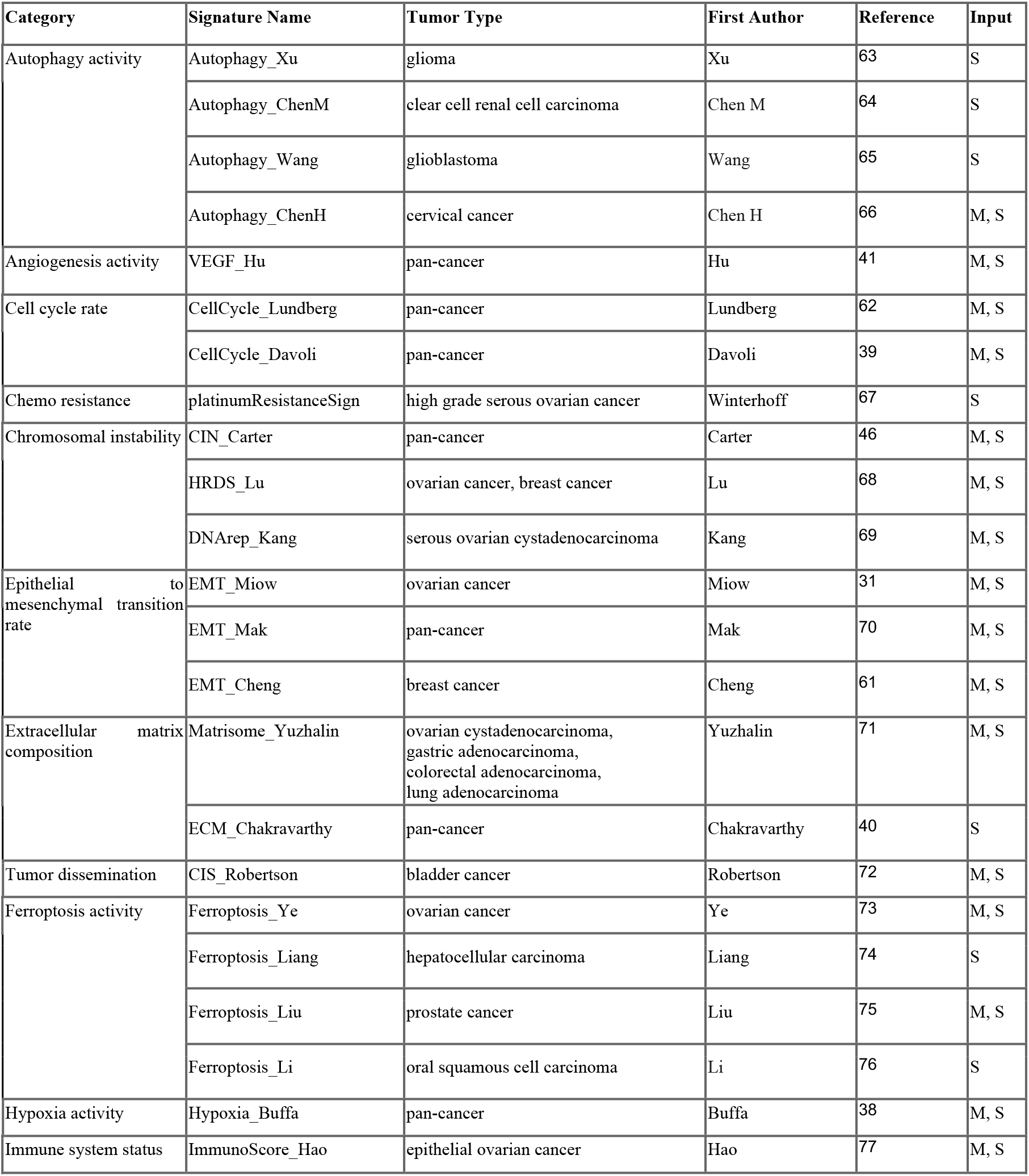
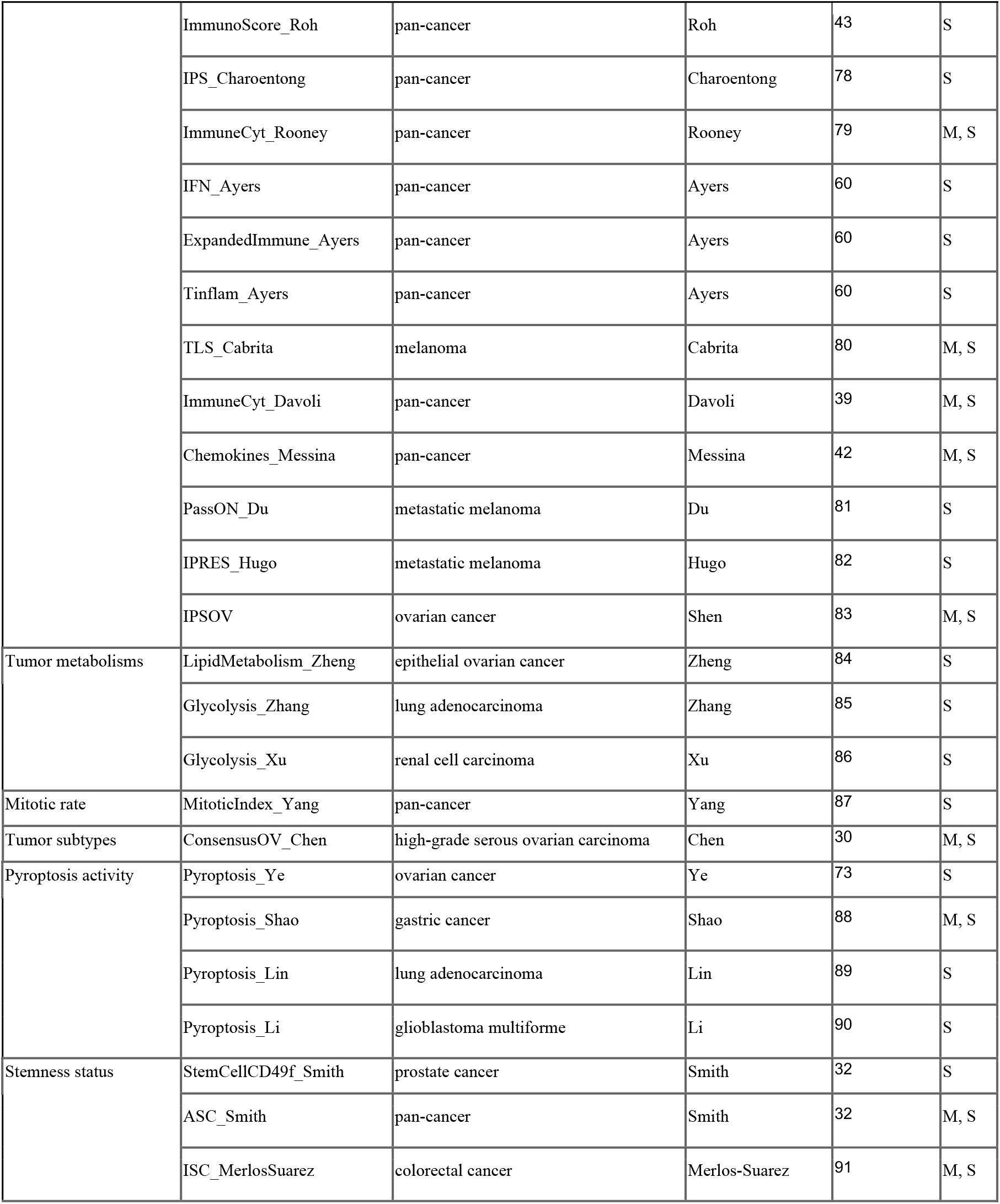
The signatures implemented in *signifinder*. The table provides the main characteristics of each signature used in *signifinder*: signature name, the tumor for which the signature was implemented, the first author of the work where the signature originated, the relevant reference, and the input types for which the signature is available (“S” for all sequencing technologies or “M” for microarray).

| Category | Signature Name | Tumor Type | First Author | Reference | Input |
| --- | --- | --- | --- | --- | --- |
| Autophagy activity | Autophagy_Xu | glioma | Xu | 63 | S |
|  | Autophagy_ChenM | clear cell renal cell carcinoma | Chen M | 64 | S |
|  | Autophagy_Wang | glioblastoma | Wang | 65 | S |
|  | Autophagy_ChenH | cervical cancer | Chen H | 66 | M, S |
| Angiogenesis activity | VEGF_Hu | pan-cancer | Hu | 41 | M, S |
| Cell cycle rate | CellCycle_Lundberg | pan-cancer | Lundberg | 62 | M, S |
|  | CellCycle_Davoli | pan-cancer | Davoli | 39 | M, S |
| Chemo resistance | platinumResistanceSign | high grade serous ovarian cancer | Winterhoff | 67 | S |
| Chromosomal instability | CIN_Carter | pan-cancer | Carter | 46 | M, S |
|  | HRDS_Lu | ovarian cancer, breast cancer | Lu | 68 | M, S |
|  | DNArep_Kang | serous ovarian cystadenocarcinoma | Kang | 69 | M, S |
| Epithelial to mesenchymal transition rate | EMT_Miow | ovarian cancer | Miow | 31 | M, S |
|  | EMT_Mak | pan-cancer | Mak | 70 | M, S |
|  | EMT_Cheng | breast cancer | Cheng | 61 | M, S |
| Extracellular matrix composition | Matrisome_Yuzhalin | ovarian cystadenocarcinoma, gastric adenocarcinoma, colorectal adenocarcinoma, lung adenocarcinoma | Yuzhalin | 71 | M, S |
|  | ECM_Chakravarthy | pan-cancer | Chakravarthy | 40 | S |
| Tumor dissemination | CIS_Robertson | bladder cancer | Robertson | 72 | M, S |
| Ferroptosis activity | Ferroptosis_Ye | ovarian cancer | Ye | 73 | M, S |
|  | Ferroptosis_Liang | hepatocellular carcinoma | Liang | 74 | S |
|  | Ferroptosis_Liu | prostate cancer | Liu | 75 | M, S |
|  | Ferroptosis_Li | oral squamous cell carcinoma | Li | 76 | S |
| Hypoxia activity | Hypoxia_Buffa | pan-cancer | Buffa | 38 | M, S |
| Immune system status | ImmunoScore_Hao | epithelial ovarian cancer | Hao | 77 | M, S |
|  | ImmunoScore_Roh | pan-cancer | Roh | 43 | S |
|  | IPS_Charoentong | pan-cancer | Charoentong | 78 | S |
|  | ImmuneCyt_Rooney | pan-cancer | Rooney | 79 | M, S |
|  | IFN_Ayers | pan-cancer | Ayers | 60 | S |
|  | ExpandedImmune_Ayers | pan-cancer | Ayers | 60 | S |
|  | Tinflam_Ayers | pan-cancer | Ayers | 60 | S |
|  | TLS_Cabrita | melanoma | Cabrita | 80 | M, S |
|  | ImmuneCyt_Davoli | pan-cancer | Davoli | 39 | M, S |
|  | Chemokines_Messina | pan-cancer | Messina | 42 | M, S |
|  | PassON_Du | metastatic melanoma | Du | 81 | S |
|  | IPRES_Hugo | metastatic melanoma | Hugo | 82 | S |
|  | IPSOV | ovarian cancer | Shen | 83 | M, S |
| Tumor metabolisms | LipidMetabolism_Zheng | epithelial ovarian cancer | Zheng | 84 | S |
|  | Glycolysis_Zhang | lung adenocarcinoma | Zhang | 85 | S |
|  | Glycolysis_Xu | renal cell carcinoma | Xu | 86 | S |
| Mitotic rate | MitoticIndex_Yang | pan-cancer | Yang | 87 | S |
| Tumor subtypes | ConsensusOV_Chen | high-grade serous ovarian carcinoma | Chen | 30 | M, S |
| Pyroptosis activity | Pyroptosis_Ye | ovarian cancer | Ye | 73 | S |
|  | Pyroptosis_Shao | gastric cancer | Shao | 88 | M, S |
|  | Pyroptosis_Lin | lung adenocarcinoma | Lin | 89 | S |
|  | Pyroptosis_Li | glioblastoma multiforme | Li | 90 | S |
| Stemness status | StemCellCD49f_Smith | prostate cancer | Smith | 32 | S |
|  | ASC_Smith | pan-cancer | Smith | 32 | M, S |
|  | ISC_MerlosSuarez | colorectal cancer | Merlos-Suarez | 91 | M, S |

### Signature implementation

Signatures are unequivocally named through a combination of the topic (or signature name) and the first author’s name (i.e., “Pyroptosis_Ye” is a signature on pyroptosis activity proposed by Ye et al.). We implemented a dedicated function for each signature, and whenever possible, signatures that rely on the same topic were grouped together. The list of available signatures and the related functions can be checked though the *signifinder* function *availableSignatures*, which lists all the needed information. Following the rules stated by the authors, *signifinder* provides its own required data for every signature (i.e., the gene list, and the corresponding coefficients and/or attributes), and takes care of the expression values input type as stated in the original work, providing automated data transformation if needed. Using *signifinder*, users can simply supply the normalized expression data to obtain signature scores. To be compatible with all forms of expression datasets – bulk, single-cell, and spatial transcriptomics – *signifinder* accepts a gene expression matrix, a data frame, a SummarizedExperiment, a SingleCellExperiment, or a SpatialExperiment object. Additionally, it manages the use of different gene IDs and expression metrics (i.e., counts, CPM, FPKM). Finally, all the signature functions return an S4 object containing the expressions - the same given by the user as input - with the addition of the signature scores inside the *colData* section.

### Signature analysis procedures and graphical summaries

Users are requested to submit normalized expression values (microarrays) or normalized counts (sequencing technologies) along with the type of data (sequencing or microarray) and the used gene ID type. *signifinder* then computes one or multiple signatures with a single command. Signatures can be selected by cancer topic, type, tissue, or a combination of these. As presented in Figure 1C, *signifinder* provides multiple ways to visually inspect the scores. Users can browse single signature plots to explore score distribution or the association with survival data. To identify the top contributor genes, users can visualize gene expression through heatmaps. Users can also compare multiple signatures to explore score distributions through ridge plots (which can also be split by user-supplied sample/cell/spot annotations), signature score heatmaps (to compare the results between samples, cells, or spots), and a correlation matrix across scores. Graphical analysis allows the detection of redundancies or specificities across signatures, which could reveal possible correlations or co-occurrences of different processes, and also allows improved patient, cell, or spot stratification.

### Signature quality check

In addition to the previously mentioned graphs, *signifinder* also provides the *evaluationSignPlot* function to assess different technical parameters that can be used to evaluate signature reliability. This was conceived to help visually find signatures that were calculated using too few genes or weak gene expressions, as well as signatures that could contain technical or biological bias.

scRNA-seq and spatial transcriptomics technologies require the detection of truly minute quantities of mRNA, which leads to the dropout of many expressed transcripts. Large percentages of indistinguishable biological and technical zero values may be detrimental for downstream signature score computation. Thus, we decided to provide a check on the total amount of counts and the percentage of zero values in each signature. Additionally, we decided it would be worth considering signatures that have scores correlated with technical or biological unwanted variability, such as batch or expression dropout, in different sample, cell, or tissue types, even though they are generally smoothed by normalization procedures^28^. Thus, for each signature, we provide the correlations of scores versus the total counts or total zero-value percentages. These parameters, in bulk RNA-seq and scRNA-seq, could highlight unwelcome results due to normalization problems and indicate the need for further assessment before making biological considerations. On the other hand, in spatial transcriptomic data, the total counts per spot reflect relevant quantitative and qualitative biological features of the tissue morphology, due to the different numbers and types of captured cells by each single spot^29^. Thus, in spatial technologies, relatively high correlations could be an intrinsic feature of the data, meaning that biological signals must be understood in context.

The *evaluationSignPlot* function returns a multi-panel plot showing for each signature: *(i)* the distribution of the sample/cell/spot average expression of the signature genes, *(ii)* the sample/cell/spot percentage of zero values in the signature genes, *(iii)* the correlation between the sample/cell/spot signature scores and total read counts (pink dots), *(iv)* the correlation between the sample/cell/spot signature scores and the percentage of total zero values (blue dots), and finally, *(v)* the percentage of signature genes used for the score calculation. This scenario can be investigated using all the samples, cells, or spots or just a subset of them. Overall, this plot allows investigators to make informed decisions about signature inclusion in downstream analyses.

### Data and code availability

*signifinder* is released under the AGPL-3 license. The source code and documents are freely available through the Bioconductor project (release 3.16). Thanks to compatibility with the *Bioconductor* data structures and procedures, *signifinder* can be used after the most popular expression data analysis packages. To enhance and simplify its usage, *signifinder* is distributed along with tutorials on how to retrieve and pre-process TCGA data, perform analysis on single or multiple signatures, and efficiently visualize results. Additionally, the code of the three case studies reported below is available in a dedicated GitHub repository (https://github.com/CaluraLab/signifinder_workflow).

### Case studies on bulk, single-cell and spatial transcriptomics

#### signifinder helps in characterizing bulk RNA-sequencing tumors by their gene expressions: a case study on a TCGA ovarian cancer dataset

Ovarian cancer, especially the most frequent and lethal high-grade serous histotype, is a disease characterized by high levels of genomic instability that impacts gene expression, causing highly variable phenotypes across patients. To demonstrate the ability of *signifinder* to automatically provide an understanding of transcriptional OVC subgroups, we analyzed 296 TCGA-OVC samples using the 11 OVC and the 17 pan-cancer signatures. Signature coverage and average expression can be found in Additional Figure 1. As an initial exploratory analysis, we computed the signature correlation matrix (Figure 2A). This helps in reducing the signatures under evaluation to highlight important ones, identifying areas of biological interest (i.e., signatures guided by the same transcriptional regulatory programs), and showing the co-occurrence of different processes.

**Figure 2.**
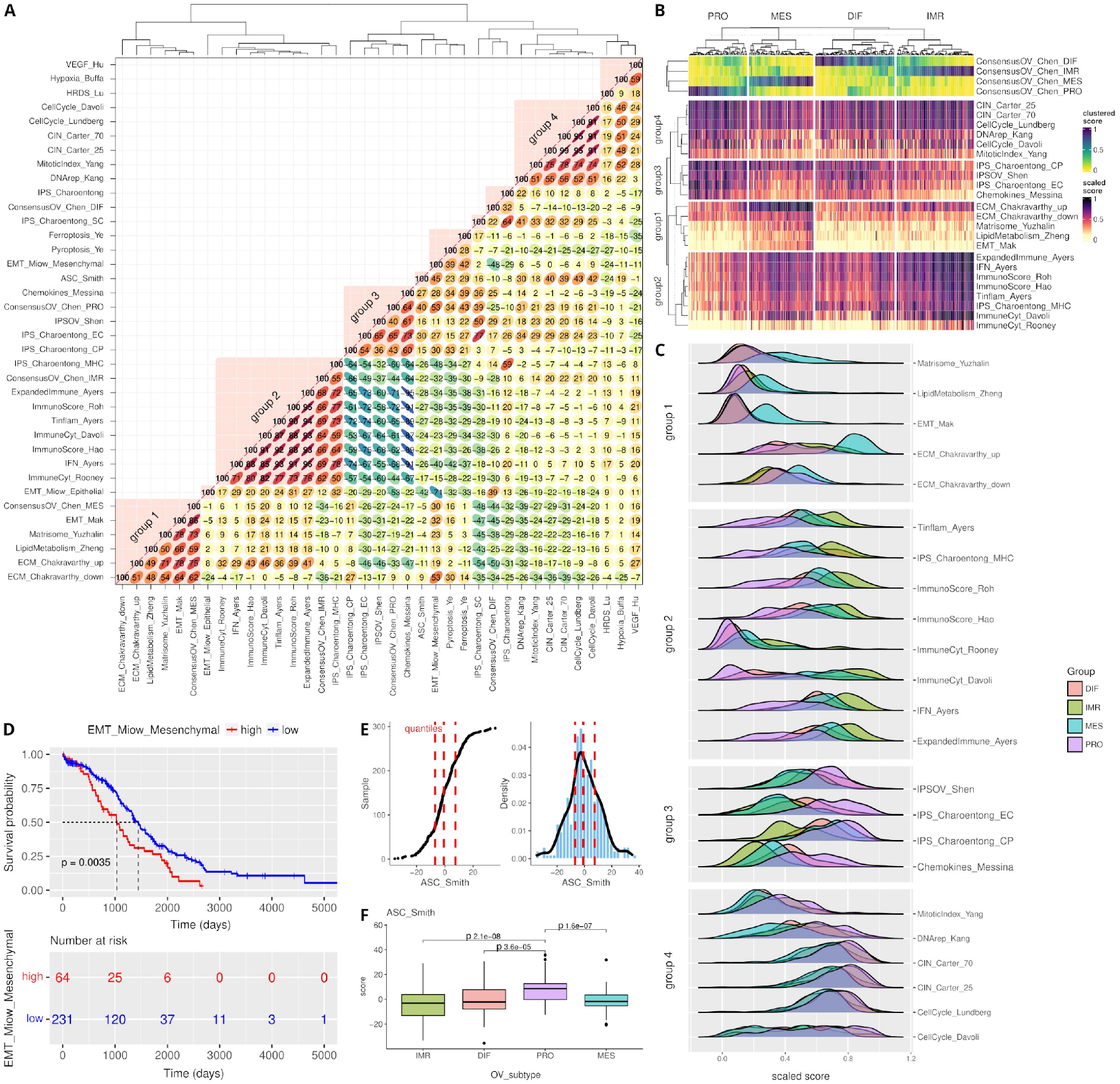
TCGA ovarian cancer dataset dissected with signifinder. **A)** Correlation matrix of pan-cancer and OVC signatures. **B)** Heatmap of signature scores of the four discussed groups of signatures with correlations with the TCGA expression subtypes, which are indicated on the top of the heatmap: PRO, IMR, MES, and DIF^30^. **C)** Ridge plot of the score distributions of the signatures in the four discussed groups. Samples are divided into the four TCGA subgroups^30^. **D)** Kaplan-Meyer plot and survival association for the EMT signature scores by Miow et al.^31^. **E)** Score distribution of the ASC signature^32^. **F)** Boxplot illustrating the ASC signature^32^ in the four TCGA transcriptional subgroups.

TCGA studies have shown that the transcriptional landscape of OVC can be differentiated into four subtypes, which are named immunoreactive (IMR), differentiated (DIF), proliferative (PRO), and mesenchymal (MES), on the basis of the gene content and previous knowledge^33^. One of the most robust and reproducible classifications of the four OVC transcriptomic subtypes is the signature *consensusOV*, which was implemented by Chen and colleagues^30^. The top panel of Figure 2B shows the four continuous scores for each sample.

The OVC signature correlation plot reported in Figure 2A shows multiple groups of signatures. The four with the highest correlations were further investigated with the plots reported in Figure 2B and 2C, where samples were stratified according to their maximum *consensusOV* score. The four groups contain three of the four major transcriptional subtypes of OVC and help to better define OVC transcriptomes.

The first group groups signatures related to ECM composition and EMT, and also includes the *consensusOV* MES score. Overexpression of genes associated with the loss of cell adhesion, developmental transcription factors, together with extracellular matrix reorganization is strongly suggestive of EMT, a process linked to poor prognosis in advanced OVC^34–36^, which is also demonstrated by the Kaplan-Meyer curve of the EMT signature in Figure 2D. The second and the third groups are composed of signatures dedicated to the many aspects of the immune system in cancer. They contain two anti-correlated behaviors: the second group (containing the *consensusOV* IMR) is characterized by signatures that collect chemokine expressions and inflammatory signals, while the third group (correlating with the *consensusOV* PRO scores) is composed of signatures associated with immune tolerance. The fourth group includes the signatures of CIN and the related mitotic index and cell cycle rate that reflect the widespread genomic alterations governing the transcriptional landscape of high-grade serous OVC.

Using the *signifinder* workflow, we described the molecular subtypes of high-grade serous OVC with an automated pipeline. This case study shows that identifying the main biological characteristics of samples is made easier by using combinations of transcriptional signatures. In addition to the description of the known TCGA-OVC molecular subtypes, *signifinder* can also reveal unexplored aspects, such as those obtained when exploring the pan-cancer signature of human Adult Stem Cells^32^ (ASC). The ASC signature, developed to mark the most aggressive epithelial cancers, shows higher scores in the OVC PRO subtype (Figure 2E-F, PRO vs *IMR p < 2.1e-08, MES p < 1.6e-07, DIFF p < 3.6e-05*). ASC are high in genes for chromosome reorganization and DNA methylation, and are considered extremely useful for pan-cancer interrogation of stem cell gene expression and for identifying additional therapeutic targets, such as the combination of treatments with DNA methyltransferase inhibitors that can sensitize tumor cells to programmed cell death^37^.

#### signifinder reveals intra-tumor cell heterogeneity: a case study of a single-cell glioblastoma sample

Glioblastoma is one of the most frequent brain tumors. Its lethality is strongly linked to tumor recurrence, mainly due to infiltrating cells that migrate from the tumor core, evading surgery and local treatment. In this case study, scRNA-seq samples from a study by Darmanis and colleagues^22^ were analyzed with the help of *signifinder*, using brain and pan-cancer signatures. Signature quality checks are shown in Additional Figure 2, for example, signatures that showed more than 90% of zeros combined with a low percentage of expressed signature genes were removed from the downstream analyses and interpretations.

In the original publication, Dermanis et al. profiled the tumor and cells from its microenvironment (i.e., the tumor core and the surrounding peripheral tissue), to reveal transcriptional and genetic variations between the two locations. Figure 3A shows a t-SNE representation of the data, color-coded by the original cell type labels as provided by the authors.

**Figure 3.**
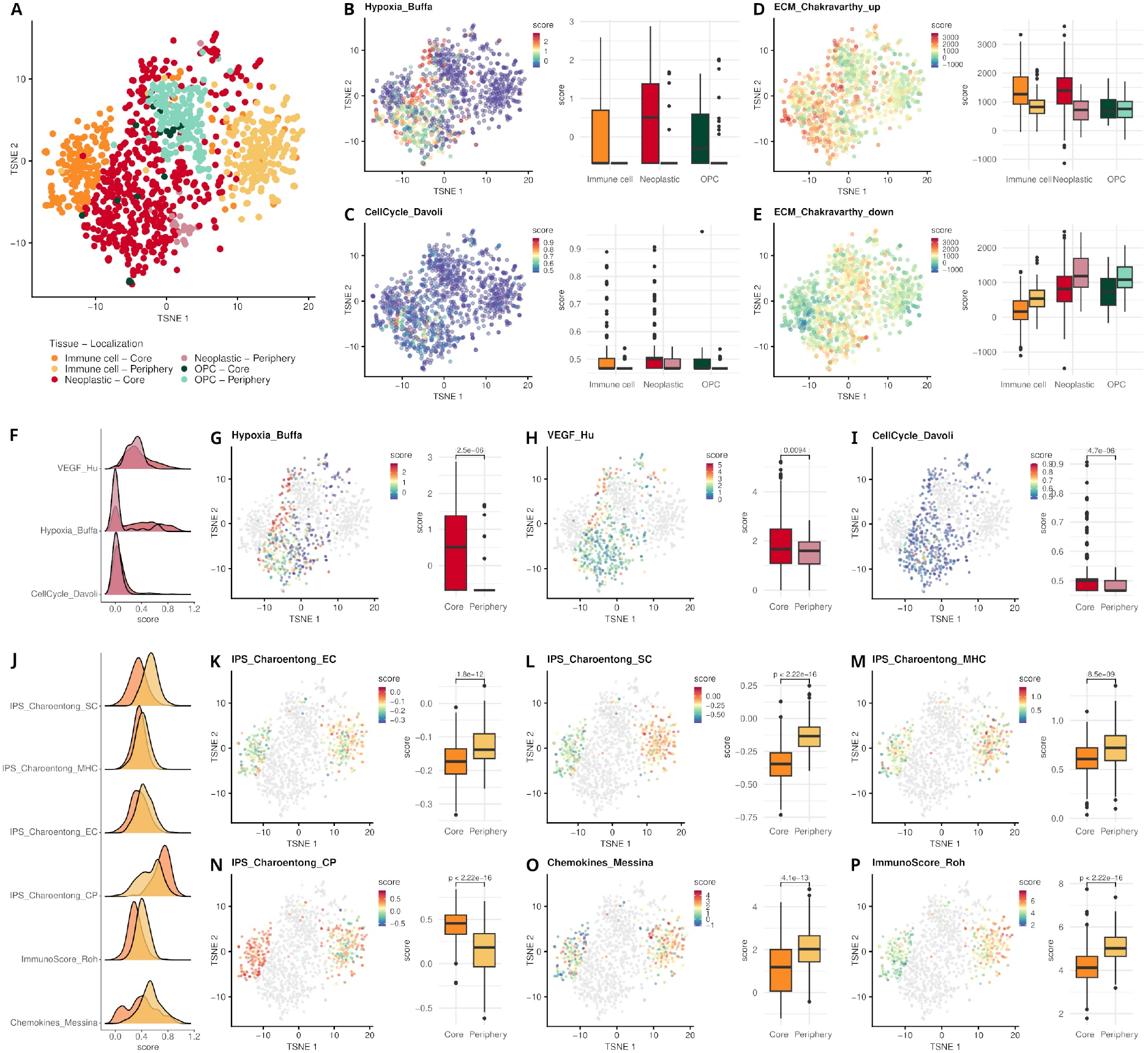
scRNA-seq data of Darmanis and colleagues^22^ studied with signifinder. **A)** t-SNE on the top 50 principal components of the expression data. Colors represent the original cell type annotations as provided by the authors additionally divided by their spatial location, i.e., tumor core or tumor periphery. Color legend is maintained throughout all the panels. t-SNE with cells colored by signature are reported in different panels together with the boxplot of signature score distributions in the different cell types and locations: **B)** Hypoxia signature scores developed by Buffa et al.^38^ and **C)** cell cycle signature scores developed by Davoli et al.^39^. **D)** ECM-Up and **E)** ECM-Down signature scores developed by Chakravarthy et al.^40^. Panels F to J are dedicated to signatures in tumor cells. **F**) Multiple signature score distributions differentiating core and peripheral tumor cells. **G)** Hypoxia signature scores developed by Buffa et al.^38^. **H)** VEGF signature scores developed by Hu et al.^41^.**I)** Cell cycle signature scores developed by Davoli et al.^30^. Panels from J to P are dedicated to signatures in immune cells. **J)** Distribution of multiple signature scores in core and peripheral immune cells. The IPS is composed of four averaged and weighted Z scores: **K)** the effector cell (EC) score, **L)** the immune-suppressive cells (SC) score, **M)** the Major Histocompatibility Complex (MHC) score, and **N)** the immune checkpoints (CP) score. **O)** The Chemokines score proposed by Messina et al.^42^, and **P)** the immune score defined by Roh et al.^43^.

*signifinder* analyses show the tumor core as a hypoxic environment for all the studied cell types, while the surrounding tissue contains cells with lower hypoxic scores, implying that they are in relatively oxygen-rich brain tissue (Figure 3B). As expected, these differences are also reported by the authors, who found hypoxia-associated angiogenesis in the tumor core compared to the periphery. The cell cycle signature defined by Davoli et al. shows that cell proliferation is confined to neoplastic cells and a small subset of immune cells, which are part of the tumor core^39^ (Figure 3C).

The ECM composition is another aspect of the TME that differentiates the transcriptional behavior of the cells in the core and the periphery. The two pan-cancer ECM signatures proposed by Chakravarty et al.^40^ (Figure 3D-E) show gradients with opposite directions between core and periphery, with neoplastic and immune cells showing higher ECM-Up scores in core cells while ECM-down is higher in peripheral cells. The ECM-up program is associated with a TGF-β-rich TME, immune evasion, and immunotherapy failure, while the ECM-down signature represents a normal-like ECM.

Focusing specifically on the neoplastic cells (Figure 3F-I and Additional Figure 3A), *signifinder* highlights their extremely heterogeneous expression profiles. Despite the significant differences between core and periphery scores, it is clear that hypoxic conditions are found only in a subgroup of neoplastic core cells (cells with high scores in Figure 3G Hypoxia_Buffa^38^,and Figure 3H VEGF_Hu signature^41^). Additionally, these hypoxic core cells also show low scores of cell cycle rate (Fig 3I, CellCycle_Davoli^39^) possibly due to the induction of cell cycle arrest in the presence of prolonged hypoxia, in order to turn off highly energy consuming processes and promote cell survival^44,45^. These hypoxic cells with low proliferation rate are spatially confined in the upper-left part of the t-SNE, which means that this condition deeply impacts the entire transcriptome of these cells. Cell proliferation is confined to a small subset of non-hypoxic cells exclusive to the tumor core, unlike peripheral tumor cells which demonstrate low proliferative potential while disseminating^39^ (Fig 3I, CellCycle_Davoli). The highly proliferative neoplastic core cells are also characterized by a high chromosomal instability score, as defined by the pan-cancer signature of Carter and colleagues^46^ (CIN_Carter_70, Additional Figure 4). This is due to the fact that genes involved in DNA replication, DNA repair, spindle assembly, and chromosome segregation include cell cycle control genes. In many tumors including gliomas, higher CIN scores are associated with unfavorable clinical outcomes and metastatic specimens. Tantalizingly, if used at the singlecell level, this evidence could be helpful in highlighting more dangerous cells for tumor progression.

Immune cells in the core and periphery are clearly separated on the t-SNE plot, showing markedly different transcriptional programs, which is also reflected by the immune signatures that report different scores in the two groups (Figure 3J and Additional Figure 3). The immunophenoscore (IPS) is composed of four averaged and weighted Z scores — the EC score that covers expression of effector cells (activated CD4^+^ T cells, activated CD8^+^ T cells, effector memory CD4^+^ T cells, and effector memory CD8^+^ T cells), the SC score that collects the expression related to immune-suppressive cells (T-regs and Myeloid-derived suppressor cells), the Major Histocompatibility Complex score for MHC-related molecules, and the immune checkpoint score to represent the activity of immune checkpoints or immunomodulators^47^. In this GB sample, all four scores showed significant differences between the core and periphery. The core seems to indicate an immunologically cold tumor, where expression of the major determinants of tumor immunogenicity is turned off (Figure 3J-N).

Another score that behaves differently between the core and periphery is the Chemokines score proposed by Messina and colleagues^42^ (Figure 3O). The score intends to predict lymphoid cell infiltrates in solid tumor masses through the expression of 12 cytokines. From the data, it seems that the presence of lymphoid cells is not homogeneous intra-mass and that most of those cytokines are expressed only by immune cells in the periphery. The pan-cancer immune score defined by Roh and colleagues is dedicated to predicting the immune checkpoint blockade treatment response. Roh and colleagues demonstrated that this immune score correlates positively with T-cell receptor clonality in pre-PD-1 blockade samples and that higher scores and T-cell receptor clonality characterize responders^43^. The Roh immune scores were found higher in peripheral cells than in core cells and thus could predict a different response of the two cell groups to PD-1 blockade treatment (Figure 3P).

#### signifinder highlights spatial-specific patterns of expression signatures: a case study on invasive ductal breast carcinoma

The spatial transcriptomic dataset presented here is a 10x Visium sample of breast invasive ductal carcinoma. Ductal carcinoma is the most common type of breast cancer (BRCA), making up nearly 80% of all BRCA diagnoses. Figure 4A shows the anatomopathologist reading of the hematoxylin and eosin staining of the FFPE sample. Multiple neoplastic areas, highlighted in red, are localized inside the ducts – this is expected in such a tumor that originates from the epithelial cells lining on the duct, and that generally, at first, invade the inner part of the ducts (carcinoma in situ), also generating necrotic areas inside the duct lumen. When the neoplastic cells overcome the duct wall, they infiltrate the stroma giving rise to the invasive carcinoma. Tumor masses are surrounded by fibrous tissue, which is delimited in blue in Figure 4A, mainly containing fibroblasts and lymphocytes. These areas are often observed in tumors like these because cancer-associated fibroblasts (CAFs) contribute to tumor proliferation through the secretion of several growth factors, cytokines, chemokines, and proteins for ECM degradation^48^. Fibrous tissue and stroma surrounding the tumor also contain areas with high densities of infiltrated lymphocytes, as indicated in Figure 4A with the light blue lines. The remainder of the section includes adipocytes and blood vessels, indicated in green and orange, respectively.

**Figure 4.**
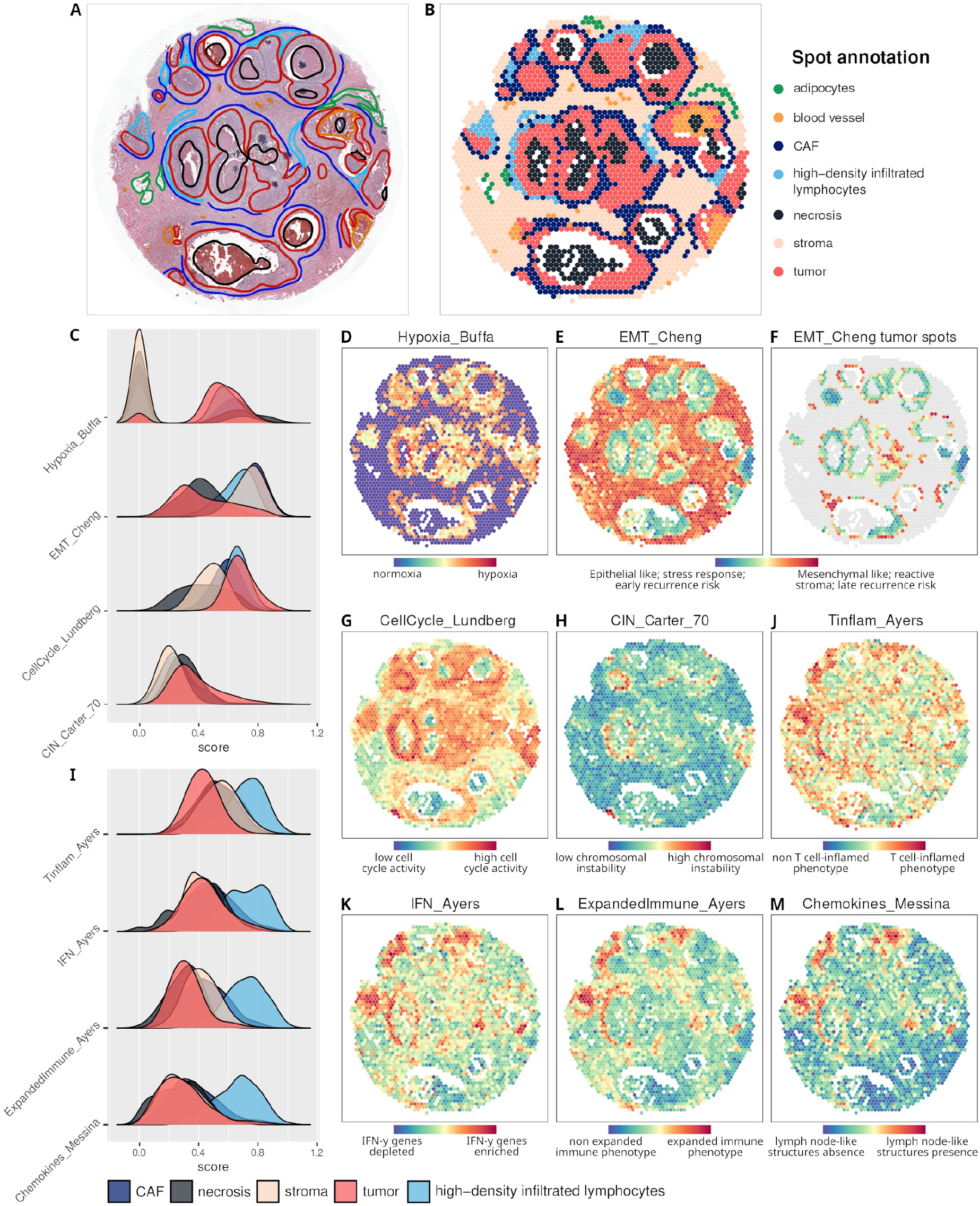
Spatial transcriptomic sample of breast invasive ductal carcinoma studied with signifinder. **A)** Histologic image with manual anatomopathological annotations. **B)** Dataset spots are annotated following manual annotations. **C)** Distribution of multiple signature scores divided into cancer associated fibroblasts (CAF), necrosis, stroma, tumor, and high-density infiltrated lymphocyte areas. Spatial score distribution of the **D)** hypoxia signature by Buffa et al.^38^, **E)** EMT by Cheng et al.^61^, **F)** EMT scores in tumor spots only, **G)** cell cycle signature by Lundberg et al.^62^, and **H)** CIN70 by Carter et al.^46^. **I)** Distribution of multiple signature scores divided into CAF, necrosis, stroma, tumor, and high-density infiltrated lymphocyte areas. Spatial score distribution of **J)** Tinflam, **K)** IFN, and **L)** Expanded Immune by Ayers et al.^60^, and **M)** Chemokines by Messina et al.^42^.

Spots are then manually classified by cell type following the anatomopathological reading of the high-resolution image (Figure 4B).

Multiple pan-cancer and BRCA signatures were then computed using the expression values of each spot, and results were compared with tissue annotations. As expected, the signatures do exhibit a relatively high percentage of zero counts as well as mild but noticeable correlations with the total count number, due to dependencies on the number of cells lying on each spot (Additional Figure 5).

The hypoxia signature by Buffa et al. shows that tumor areas are highly hypoxic compared to the normal stroma^38^ (Figure 4C and D), and that all cell types show high variance of hypoxia scores (Figure 4C), with the highest scores in certain necrotic spots.

The tumor specimens are characterized by the coexistence of both epithelial and mesenchymal markers with a particular spatial distribution. The EMT signature by Cheng and colleagues, which was originally provided as a prognostic biomarker associated with late-disease recurrence in BRCA, is made of 51 genes that are mainly involved in EMT. High scores are localized to the leading edge of tumors in close apposition with the surrounding stroma. Starting from the duct basement membrane, where the tumor arises, the score describes a decreasing pattern when moving to the inner part of the duct (Figure 4C, E, and F). Interestingly, since high EMT scores are associated with high-risk of late recurrence, the spatial distribution of this score suggests that the hyperproliferative cells found in the basement membrane (i.e., the origin cells of these tumors) are also the most dangerous cells for relapse. Tumor spots show high cell-cycle rates, as determined by the Lundberg et al. signature that lights up the tumor and the nearby areas (Figure 4C, G). The lowest scores remain confined to the non-reactive stroma and to necrotic areas. Thus, we can appreciate here that tumor cells show heterogenic proliferation rates. Interestingly, the highest proliferative tumor areas also have high levels of CIN (Figure 4C, H) as shown by the CIN signature proposed by Carter et al., which is based on expression aberrations in genes localized to each chromosomal region^46^. In ductal breast carcinoma, the presence of a high rate of chromosomal copy number variations is associated with an increased immune response, as well as the presence of tumor-infiltrating lymphocytes and PD-L1 gene expression^49–52^. We can confirm that high-CIN tumor areas are in close proximity with those reported to have high lymphocyte density.

In recent years, it has been generally accepted that these immune-related areas play a role as prognostic and predictive markers in invasive BRCA^53–56^. However, some controversial evidence suggests that their predictive value depends on the exact composition of the infiltrate^57–59^, which is almost impossible to assess based solely on hematoxylin and eosin staining. Signatures can thus help in evaluating the immune activities of these zones. Multiple immune system signatures characterize the areas with high lymphocyte density: Tinflam, IFN (interferons), and Expanded Immune by Ayers et al.^60^, as well as the Chemokines signature proposed by Messina et al.^42^. These signatures show the highest values in proximity to areas with high densities of lymphocytes (Figure 4J-M, and Additional Figure 6). The signatures from Ayers and colleagues were developed to identify tumors with a T cell-inflamed microenvironment, characterized by active IFN-γ signaling, cytotoxic effector molecules, antigen presentation, and T cell active cytokines, which are common features of tumors that are responsive to PD-1 checkpoint blockade. Similarly, the Chemokines signature, which was independently developed with similar purposes to identify tumor-localized ectopic lymph node-like structures, shows high scores in the same areas^42^.

## Conclusions

The release of a computational algorithm along with key cancer signatures would be a major step towards a future of assured reproducibility and usability. However, to date, this concept has received little attention. By developing *signifinder*, we have achieved three main results: *(i)* we have created the infrastructure to collect and implement transcriptomic signatures of diverse origin (bulk, single-cell, and spatial transcriptomic data); *(ii)* we have provided a compendium of cancer gene signature implementations, by collecting and screening papers from the literature; and, (iii) by allowing single-cell and spatial transcriptomic datasets to be used as inputs, we have provided a tool that can test how bulk gene expression signatures behave intra-tumor, thus finally investigating the ability of these signatures to detect different cancer cell states.

This project is one of the first attempts at providing a computational infrastructure for the systematic investigation of gene expression signatures in cancer. Indeed, because *signifinder* is implemented in R, it is compatible with most of the most common tools and pipelines for transcriptome data analysis. *signifinder* is also built with additional functions to provide easier visualization and interpretation of the results, thus simplifying the understanding of the role that multiple hallmarks within and between patients, samples, cells, and tissues can play.

Moving forward, a general agreement on how molecular subgroups of cancer are technically defined would facilitate the use of expression data in clinical management. *signifinder* easily allows comparison across multiple signatures, highlighting processes, interactions, and cooperation. It also helps to determine cancer cell states in cell-level transcriptomes. A better understanding of the biology underlying these subgroups and their cell-level heterogeneity will provide more rational assessments and targeted treatments.

Future directions of the *signifinder* package involve continuous addition and implementation of new signatures. The use of *signifinder* on new high-resolution technologies, such as singlecell RNA-seq and spatial transcriptomics, is perhaps the most promising way of putting *signifinder* to work. Cancer cells exploit existing gene expression modules, expressing them at different levels and with different levels of heterogeneity. With this tool, heterogeneity in these systems can be finally evaluated, quantified, and weighted to evaluate the importance of the signature prediction at the patient level. This means that *signifinder* could pave the way for the eventual automatic assessment of cancer cell states in high-throughput expression datasets and have a role in prediction of treatment options.

## Data and code availability

The source code and documents have been released through the Bioconductor platform (https://bioconductor.org/packages/release/bioc/html/signifinder.html). The developing version of the package is also available at https://github.com/CaluraLab/signifinder. To assure reproducibility, the code that was used for the three case studies is also available in a dedicated GitHub repository (https://github.com/CaluraLab/signifinder_workflow).

## Additional File

- Additional Figure 1 - Signature evaluation plot for the RNA-seq TCGA OVC dataset.
- Additional Figure 2 - Signature evaluation plot for the GB sample of scRNA-seq data by Darmanis and colleagues^22^.
- Additional Table 1 - The number of cells from patient BT_S2 from the dataset of Darmanis and colleagues^22^ divided by cell location (core or periphery) and cell type.
- Additional Figure 3 - Correlation plots of selected signatures in (A) tumor and (B) immune cells from the dataset of Darmanis and colleagues^22^.
- Additional Figure 4 - Scatterplot of CellCycle_Davoli scores and CIN_Carter_70 scores computed on neoplastic cells from the Darmanis et al.^22^ GB scRNA-seq dataset.
- Additional Figure 5 - Signature evaluation plot for the Visium 10x sample of breast invasive ductal carcinoma.
- Additional Figure 6 - Spatial score distribution of Tinflam_Ayers, IFN_Ayers, ExpandedImmune_Ayers, and Chemokines_Messina for the spots annotated as “high-density infiltrated lymphocytes”, “CAF”, and “tumor”.
- Additional Figure 7 - Heatmap of the log2 expression values of genes composing the Tinflam_Ayers, IFN_Ayers, ExpandedImmune_Ayers, and Chemokines_Messina signatures in the spatial transcriptomics ductal BRCA case study.

## Abbreviations

Abbreviations: Meaning

ASC: Adult Stem Cell

BRCA: Breast Cancer

CAFs: Cancer Associated Fibroblasts

CIN: Chromosomal Instability

DIF: Differentiated

ECM: Extracellular Matrix

EMT: Epithelial-to-Mesenchymal Transition

FFPE: Formalin-Fixed Paraffin-Embedded

GB: Glioblastoma

IFN: Interferons

IMR: Immunoreactive

IPS: Immunophenoscore

MES: Mesenchymal

MHC: Major Histocompatibility Complex

OVC: Ovarian Cancer

PRO: Proliferative

scRNA-seq: Single cell RNA Sequencing

t-SNE: t-Distributed Stochastic Neighbor Embedding

TCGA: The Cancer Genome Atlas

TME: Tumor Microenvironment

## Author contributions

SP developed the signifinder package and coordinated contributions to it. SP and AC performed the data analyses, LM contributed to the documentation and vignette of the package and FP contributed to signature collection and implementation. LM extensively tested the analysis and package code. GE provided the histopathological image reading of the 10x spatial transcriptomic data. PM and DR helped in package development, and PM, DR and CR contributed to conceive the signature evaluation strategies. EC and SP drafted the manuscript, PM, CR, DR and LM reviewed and edited the manuscript. EC and CR conceived the project, and EC supervised the project. All Authors read and approved the final manuscript.

