## Additional for "signifinder enables the identification of tumor cell states and cancer expression signatures in bulk, single-cell and spatial transcriptomic data"

### Additional Figure 1

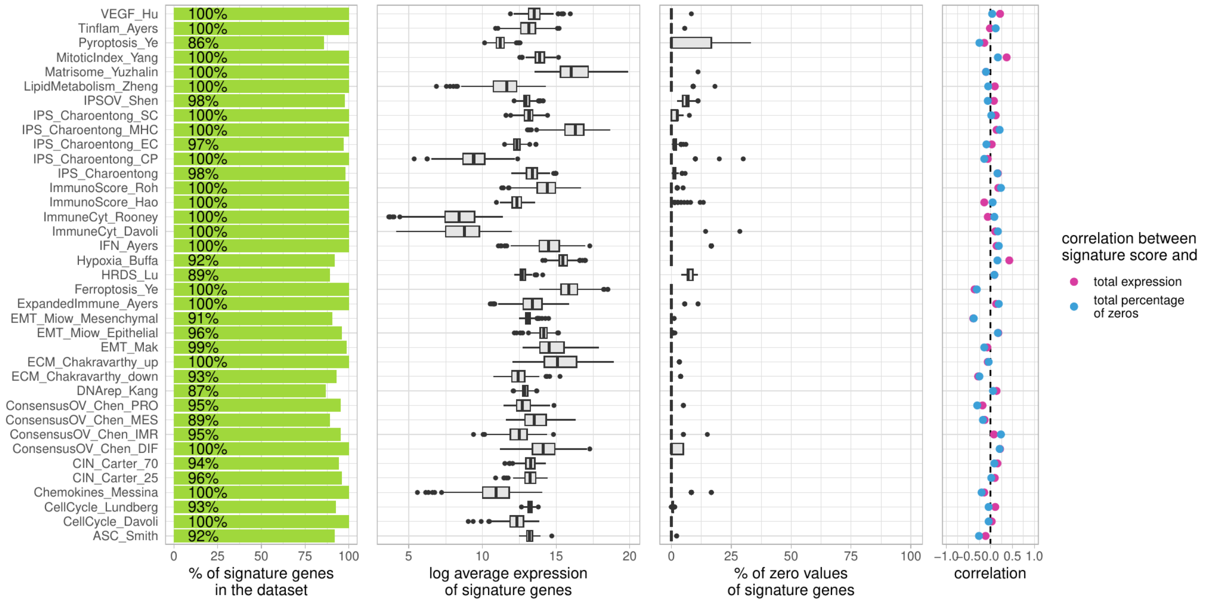


*Signature evaluation plot for the bulk ovarian cancer case study.*

### Additional Figure 2

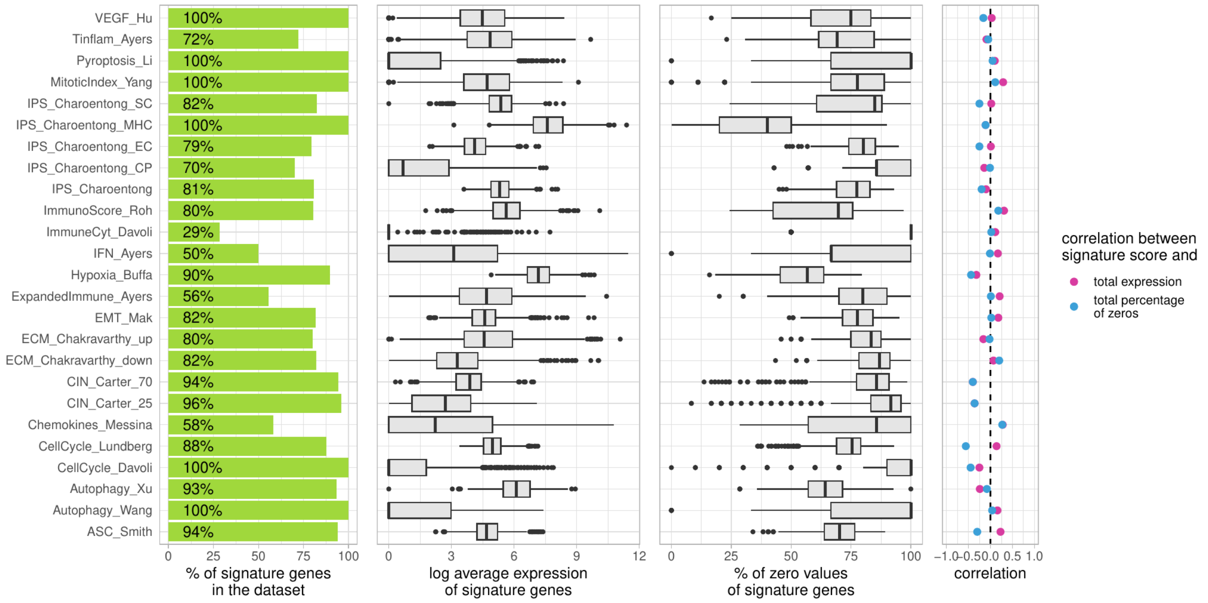


*Signature evaluation plot* for the single-cell glioblastoma case study

### Additional Table 1

| | Astrocyte | Immune cell | Neoplastic | Neuron | Oligodendrocyte | OPC | Vascular |
|---|---|---|---|---|---|---|---|
| Periphery | 26 | 200 | 27 | 17 | 15 | 154 | 0 |
| Tumor core | 0 | 184 | 468 | 1 | 4 | 21 | 1 |

*The table contains the original number of cells from patient BT_S2, divided by the cell location and the cell type. In order to have sizable cohorts for signature score comparisons, cells were filtered by type. Following the authors original cell type annotations, we kept only those present in both the tumor core and the tumor peripheral samples with a sample size greater than 20. Thus, immune cells, neoplastic and oligodendrocyte precursor cells (OPC) were considered for further analysis.*

### Additional Figure 3

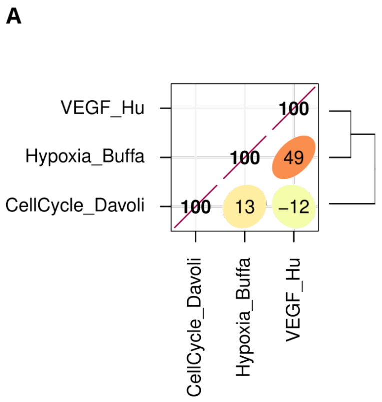


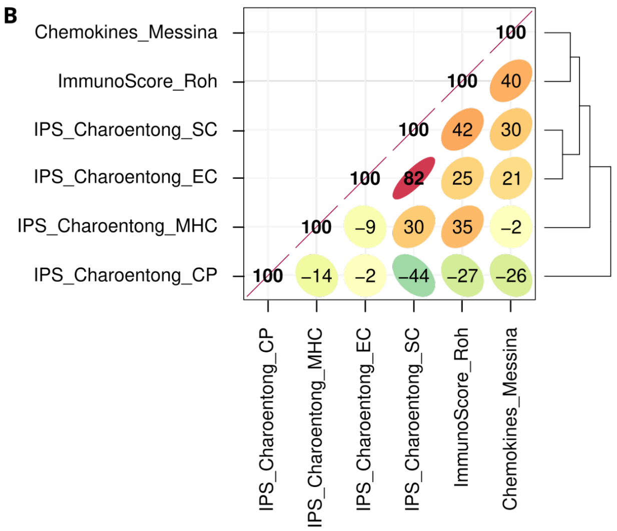


Correlation matrices on glioblastoma single-cell dataset. **A)** Correlation between signature scores of VEGF_Hu, Hypoxia_Buffa and CellCycle_Davoli computed on the neoplastic cells. **B)** Correlation between signature scores of Chemokines_Messina, ImmunoScore_Roh and IPS_Charoentong computed on the immune cells.

### Additional Figure 4

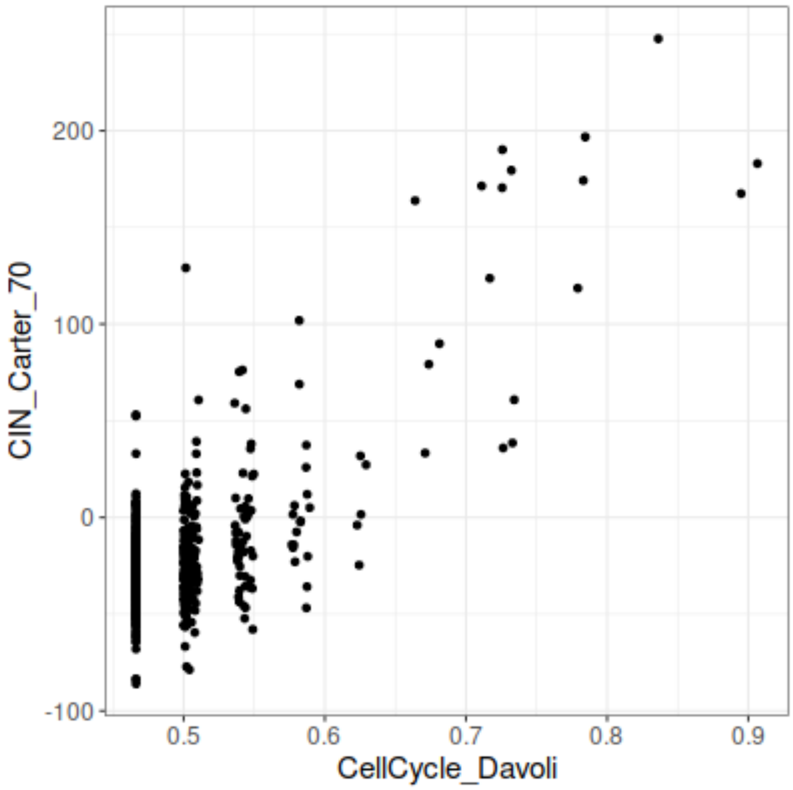


Scatterplot of CellCycle_Davoli scores and CIN_Carter_70 scores computed on the neoplastic cells from the Darmanis et al. glioblastoma single-cell dataset.

### Additional Figure 5

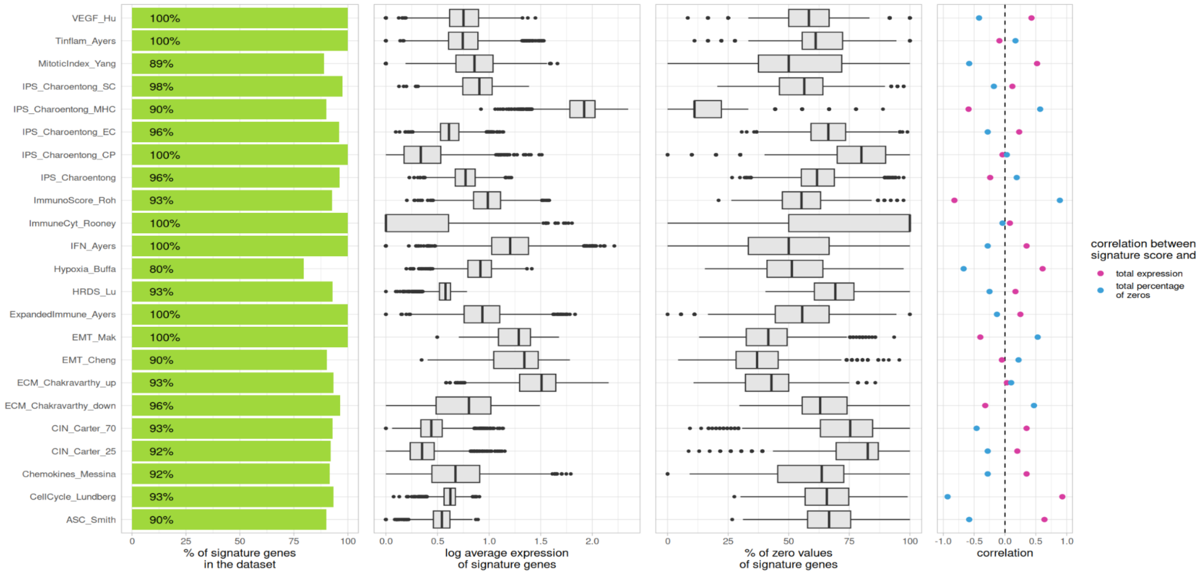


*Signature evaluation plot for the spatial breast cancer case study.*

**Additional Figure 6**

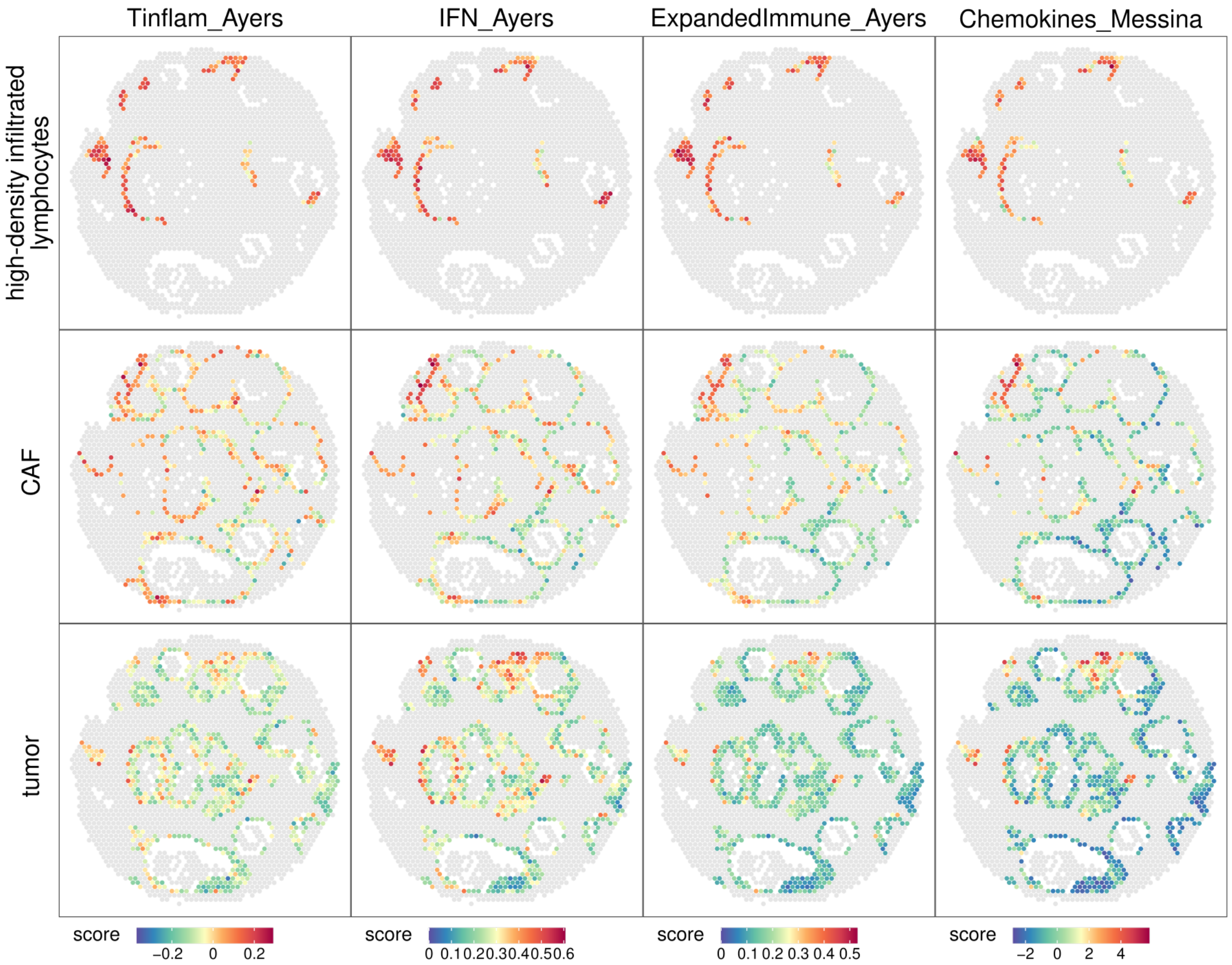


*Spatial score distribution of Tinflam_Ayers, IFN_Ayers, ExpandedImmune_Ayers and Chemokines_Messina for the spots annotated as "high-density infiltrated lymphocytes", "CAF" and "tumor".*

### Additional Figure 7

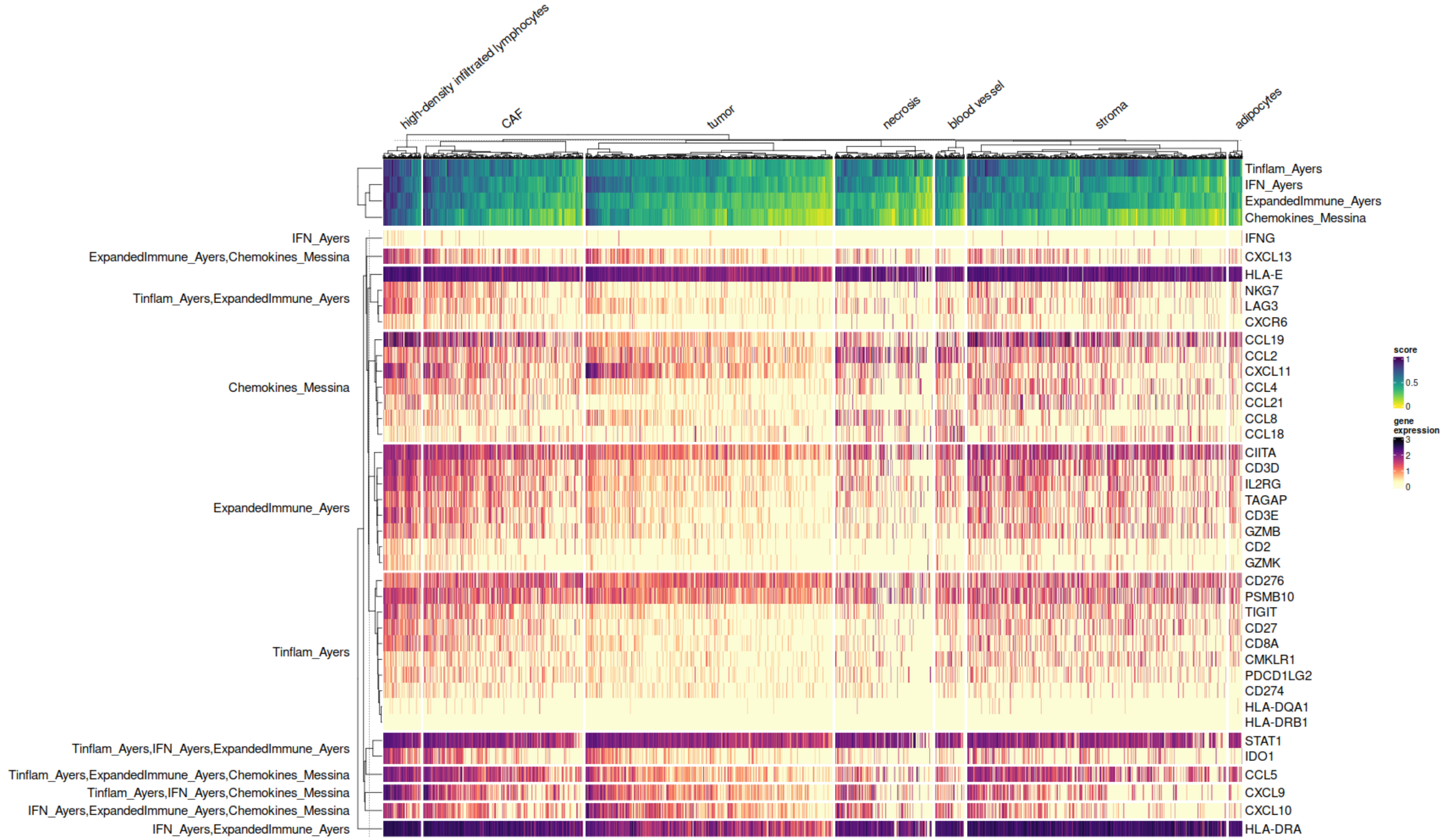


*Heatmap of the log2 expression values of genes composing the Tinflam_Ayers, IFN_Ayers, ExpandedImmune_Ayers and Chemokines_Messina signatures in the spatial transcriptomics ductal breast cancer case study.* Genes in rows are grouped by signatures and spots in columns are grouped by anatomopathological annotations.
